# cfDNAanalyzer: a comprehensive toolkit for analyzing cell-free DNA genomic sequencing data in liquid biopsy

**DOI:** 10.1101/2025.06.09.658767

**Authors:** Junpeng Zhou, Keyao Zhu, Xiaoqian Huang, Jinqi Yuan, Yumei Li

## Abstract

Liquid biopsy, through the analysis of plasma cell-free DNA (cfDNA), is transforming diagnostic medicine by enabling non-invasive insights into the genomic and epigenomic landscape of diseases—most notably cancer. As cfDNA reflects dynamic contributions from multiple tissues, it holds immense potential for early detection, disease monitoring, and personalized treatment. However, the rapid expansion of cfDNA applications has outpaced the availability of unified, versatile computational tools for the systematic analysis of cfDNA data. To address this unmet need, we present cfDNAanalyzer—a comprehensive, user-friendly toolkit that streamlines the feature extraction, selection, and construction of machine learning-based models for disease detection using cfDNA. Designed for flexibility and scalability, cfDNAanalyzer supports multimodal feature extraction, integration, and interpretable outputs, enabling both exploratory research and translational applications. By standardizing and accelerating cfDNA data analysis, cfDNAanalyzer provides a critical resource to advance precision diagnostics and deepen biological insight into disease processes.

## Introduction

The advent of liquid biopsy has revolutionized the field of personalized medicine, offering a minimally invasive method for detecting and monitoring a range of diseases, particularly cancer. Cell-free DNA (cfDNA), released into bodily fluids by both healthy and diseased cells^1–3^, carries valuable genomic and epigenomic information that can be leveraged for early disease detection and real-time monitoring of disease progression. One widely used approach is the genomic sequencing of cfDNA, providing critical insights into genetic alterations, such as copy number variations^4,5^. However, identifying disease-specific genetic changes is challenging due to the low concentration of mutated DNA fragments in patient samples and the limited number of detectable genetic alterations^6–9^.

In recent years, the field of cfDNA analysis has expanded beyond traditional genetic profiling. cfDNA is released into the bloodstream through apoptotic and necrotic events, during which endonucleases fragment DNA. This process creates distinct fragmentation patterns where closed chromatin regions, protected by nucleosomes, remain intact, while open chromatin regions are more sensitive to endonuclease activity^2^. These fragmentation signatures carry epigenetic information that can reveal the tissue of origin, chromatin structure, and nucleosomal organization^2,10^. To date, various fragmentomics features derived from cfDNA sequencing data have emerged as promising biomarkers for disease detection. These include fragmentation profile^11^, end motif frequency and motif diversity score^12,13^, windowed protection score (WPS)^2^, orientation-aware cfDNA fragmentation (OCF) features^10^, nucleosome profile^14^, nucleosome occupancy and fuzziness^15^. Furthermore, cfDNA fragmentation patterns around transcription start sites (TSS), such as promoter fragmentation entropy (PFE) and coverage profile, have been utilized to infer gene expression states, offering new opportunities for cancer detection and tissue origin inference^16,17^.

Despite the rapid development of fragmentomics, the current bioinformatics tools for cfDNA feature extraction remain inadequate. They are often not standalone but rather embedded within workflows that are not user-friendly. To date, only three major cfDNA analysis toolkits have been published, each with notable limitations^18–20^. These existing toolkits are limited in their ability to comprehensively extract a wide range of fragmentomics features, often restricting users to a narrow subset of biomarkers. Moreover, they require multiple manual steps for all analyses, which is not user-friendly. Perhaps most importantly, none of these toolkits offer direct and complete capabilities for downstream processing, such as feature processing and selection, as well as building machine learning models that can incorporate diverse cfDNA features for robust disease detection and classification (**Supplementary Table 1**). Given the increasing reliance on liquid biopsy for biomarker discovery, there is an urgent need for a more flexible, comprehensive toolkit that can streamline the analysis of cfDNA data while enabling cutting-edge machine learning applications.

To address these gaps, we present **cfDNAanalyzer** (<u>c</u>ell-<u>f</u>ree <u>DNA</u> sequencing data <u>analyzer</u>), a comprehensive and user-friendly toolkit specifically designed for the analysis of cfDNA genomic sequencing data. This all-in-one solution enables feature extraction, processing, selection, and the integrated analysis of multiple cfDNA genomic and fragmentomics signatures, culminating in machine learning-based disease classification. By unifying these functionalities within a single framework, cfDNAanalyzer streamlines and standardizes the interpretation of liquid biopsy data, empowering researchers and clinicians to accelerate the discovery and validation of novel biomarkers for disease detection and monitoring.

## Results

### cfDNAanalyzer: a comprehensive toolkit for cfDNA genomic sequencing data analysis

The cfDNAanalyzer toolkit is organized into three core modules, designed to facilitate the comprehensive analysis of cfDNA genomic sequencing data: (1) *Feature Extraction*, which extracts genomic and fragmentomic features at whole-genome or region-specific levels; (2) *Feature Processing and Selection*, enabling refinement and optimization of extracted features through various statistical and machine learning methods; and (3) *Machine Learning Model Building*, supporting the development of predictive models for disease detection and classification (**Fig. 1**). To ensure seamless analysis, cfDNAanalyzer also provides scripts for converting raw sequencing data into compatible formats, along with tutorials for data visualization—delivering a streamlined, command-line-based, all-in-one solution for cfDNA research.

**Fig. 1.**
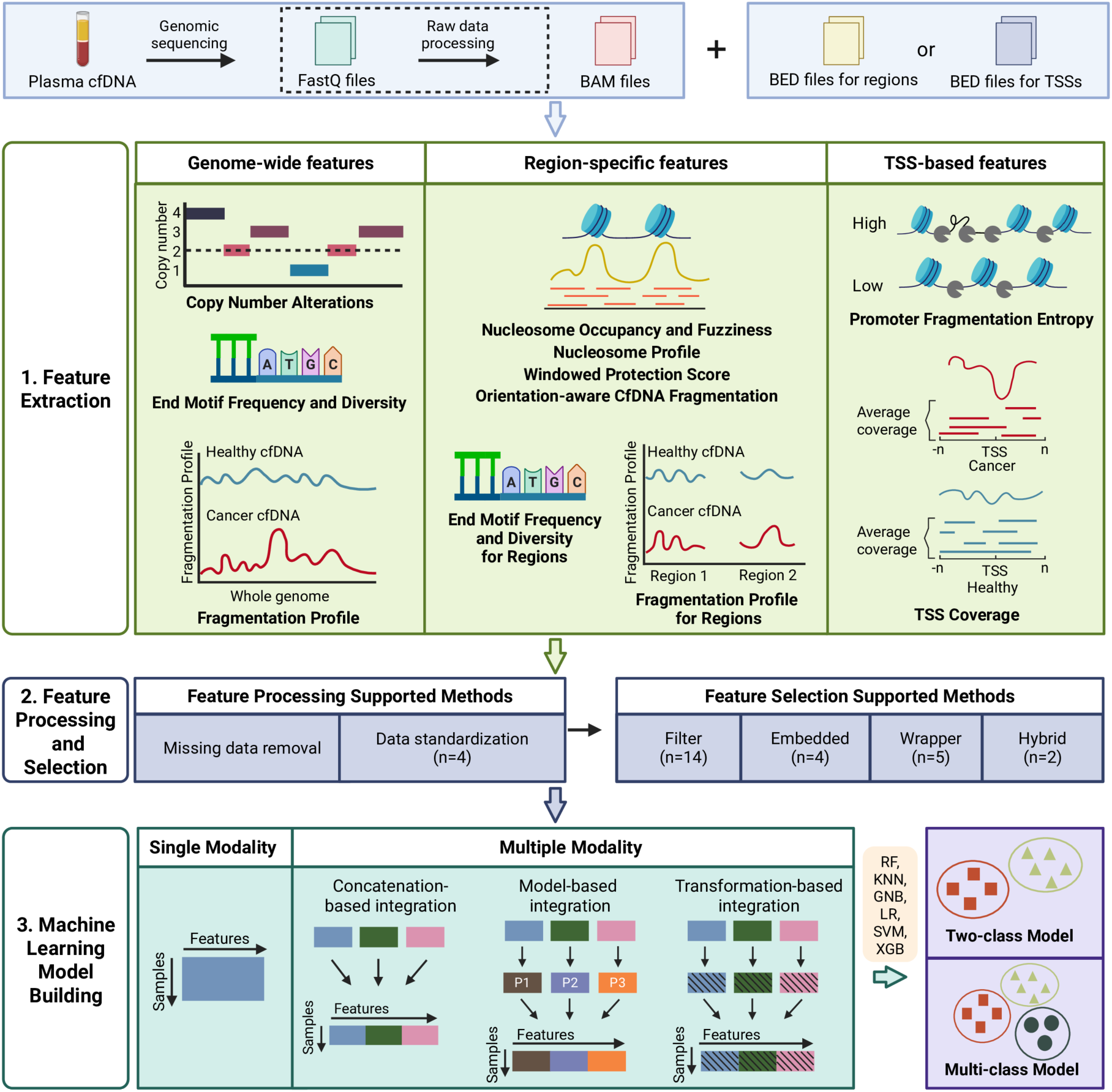
Workflow of cfDNAanalyzer. The pipeline comprises three core components: Feature Extraction, Feature Processing and Selection, and Machine Learning Model Building. The schematic illustrates the data flow between components and highlights the key functionalities within each stage, from raw cfDNA sequencing data to model-based classification or prediction.

The workflow begins with input preparation, including cfDNA sequencing data in BAM format and optional BED files for region-specific analyses. To simplify preprocessing, a utility script is provided for converting raw FASTQ files to BAM format. The first core module, *Feature Extraction*, supports a wide range of genomic and fragmentomic features that can be analyzed across the whole genome, specific regions, or TSS. Where applicable, cfDNAanalyzer incorporates source code from original studies to ensure methodological consistency. It is noteworthy that cfDNAanalyzer is capable of processing multiple samples and features simultaneously, making it an efficient solution for analyzing large clinical datasets. The extracted feature matrices are then passed to the *Feature Processing and Selection* module, which handles missing data, standardizes the data, and applies a wide range of feature selection techniques, including filter, wrapper, embedded, and hybrid approaches (**Supplementary Table 2**). The final module, *Machine Learning Model Building,* enables the construction of predictive models using individual features or integrated multi-feature approaches to enhance predictive power. With support for both binary (e.g., disease vs. normal) and multi-class (e.g., cancer subtypes) classification tasks, the toolkit is versatile enough to handle a variety of research and clinical applications. Moreover, the ability to evaluate different feature integration strategies—such as concatenation-based, model-based, and transformation-based approaches—offers users flexibility in designing models tailored to their specific datasets and research questions (**Supplementary Table 3**).

All steps, from data preparation to model development, are executed via a single streamlined command (*cfDNAanalyzer.sh*), providing a seamless, end-to-end solution. This one-command-line design significantly lowers the barrier to cfDNA analysis, making the toolkit accessible to both novice users and experienced bioinformaticians.

### Comprehensive exploration of features extracted by cfDNAanalyzer

To demonstrate the utility of cfDNAanalyzer’s feature extraction module, we applied it to a whole-genome sequencing (WGS) dataset comprising multiple cancer types and healthy controls (multi-cancer dataset; **Supplementary Table 4**). This evaluation simulates a typical application scenario, where users aim to comprehensively explore cfDNA characteristics for biomarker discovery using the toolkit’s built-in tutorial.

cfDNAanalyzer efficiently extracted a wide array of genomic and fragmentomic features. For instance, copy number alternations (CNA) and fragmentation profiles were visualized as chromosome-level tracks, with color-coded samples enabling clear comparison between cancer patients and healthy controls (**Fig. 2A-B; Supplementary** Fig. 1A-B). Distinct CNA patterns and fragmentation signatures characteristic of cancer were readily observed. Additionally, we compared end motif frequencies across the entire genome, highlighting top motifs with the most significant differences between cancer and control samples (**Fig. 2C; Supplementary** Fig. 1C). These visualizations provide immediate insights into cfDNA alterations, making it easier to discern patterns that may be indicative of disease progression.

**Fig. 2.**
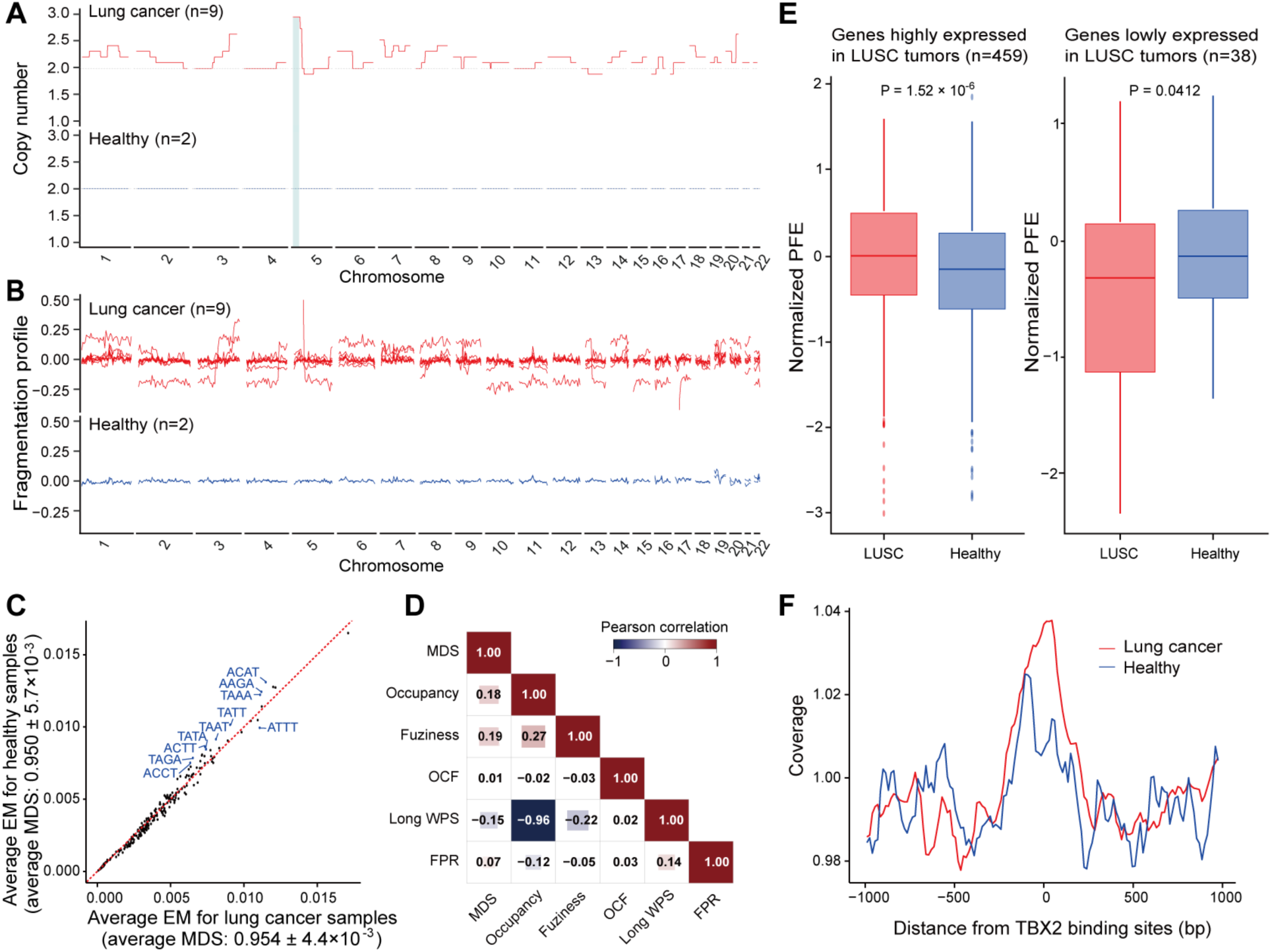
Comparative analysis of cfDNA features between lung cancer patients and healthy controls. **(A)** Copy Number Alteration (CNA) profiles on autosomes for lung cancer samples (n=9) versus healthy controls (n=2). Genomic regions with >1.5-fold CNA differences are highlighted by light blue rectangles. **(B)** Differential Fragmentation Profile (FP) patterns across autosomes, showing differential distributions between lung cancer and healthy samples. **(C)** Comparison of average End Motif frequency and diversity (EM), with the 10 most significant altered motifs highlighted in blue. **(D)** Pearson correlation matrix of fragmentomic features for ±1000 bp regions around TSS in healthy samples. **(E)** Normalized Promoter Fragmentation Entropy (PFE) differences in lung squamous cell carcinoma (LUSC) and healthy samples, stratified by gene expression levels in LUSC tumors versus normal tissues: highly expressed genes (left) and lowly expressed genes (right), with the gene numbers indicated accordingly. **(F)** Aggregative lines showing cfDNA coverage at TBX2 binding sites (±1000 bp) in lung cancer and healthy samples.

At the regional level, the toolkit enables the exploration of feature relationships to guide downstream machine learning. For instance, in promoter regions, nucleosome occupancy and WPS exhibited a strong negative correlation in the analyzed dataset (Pearson correlation coefficient = – 0.96; **Fig. 2D; Supplementary** Fig. 2), suggesting potential redundancy. In contrast, OCF showed a weak correlation with other features, highlighting its complementary value. Such analyses help users prioritize non-redundant, informative features for model development. For TSS-centered features, users can assess whether PFE significantly differs between cancer and healthy samples, particularly for genes known to be differentially expressed (**Fig. 2E; Supplementary** Fig. 1D). Moreover, cfDNAanalyzer supports extraction and comparison of nucleosome profiles across 377 transcription factor binding site (TFBS) sets or user-defined regions, uncovering cancer-specific patterns of nucleosome organization (**Fig. 2F; Supplementary** Fig. 3). This region-specific analysis not only facilitates informed feature selection for modeling but also deepens our understanding of regulatory alterations in disease.

In summary, the exploration of features extracted by cfDNAanalyzer not only streamlines the process of identifying the most relevant features for model construction but also enhances biological interpretation. The toolkit’s comprehensive feature set and intuitive visualization tutorial empower researchers to make informed decisions, facilitating both the discovery of novel biomarkers and the study of cancer progression. By combining efficiency, flexibility, and depth, cfDNAanalyzer significantly advances the analysis of cfDNA in liquid biopsy research.

### Optimized feature selection and functional interpretation

Following initial feature exploration, cfDNAanalyzer was employed to identify the most informative features using 25 feature selection methods spanning four major categories: filter, embedded, wrapper, and hybrid approaches. To benchmark performance, we applied these methods to both the multi-cancer dataset and a newly collected WGS dataset comprising 25 breast cancer patients and 25 healthy controls (breast cancer dataset; **Supplementary Table 4**). We compared the results of principal component analysis (PCA) before and after feature selection to guide the choice of feature selection method.

Among the tested methods, Random Forest importance, Boruta, Fast Correlation-Based Filter, and Fisher’s Score demonstrated notable improvement in at least one of the datasets (**Supplementary** Fig. 4A**; Supplementary** Fig. 5A), as evidenced by a higher proportion of variance captured by the top three principal components (PCs) and improved class separation in PCA plots (**Fig. 3A-B; Supplementary** Fig. 4B-E**; Supplementary** Fig. 5B-C). Further supporting their utility, functional enrichment analysis confirmed the biological relevance of the selected features (**Fig. 3C and Supplementary** Fig. 5D). For instance, the enrichment of the JAK-STAT signaling pathway, known to play a key role in cancer progression^21–23^, highlights their potential as biomarkers for cancer detection and disease monitoring. Together, PCA-based evaluation and functional annotation help users identify robust, biologically meaningful features, streamlining the refinement process prior to machine learning model development.

**Fig. 3.**
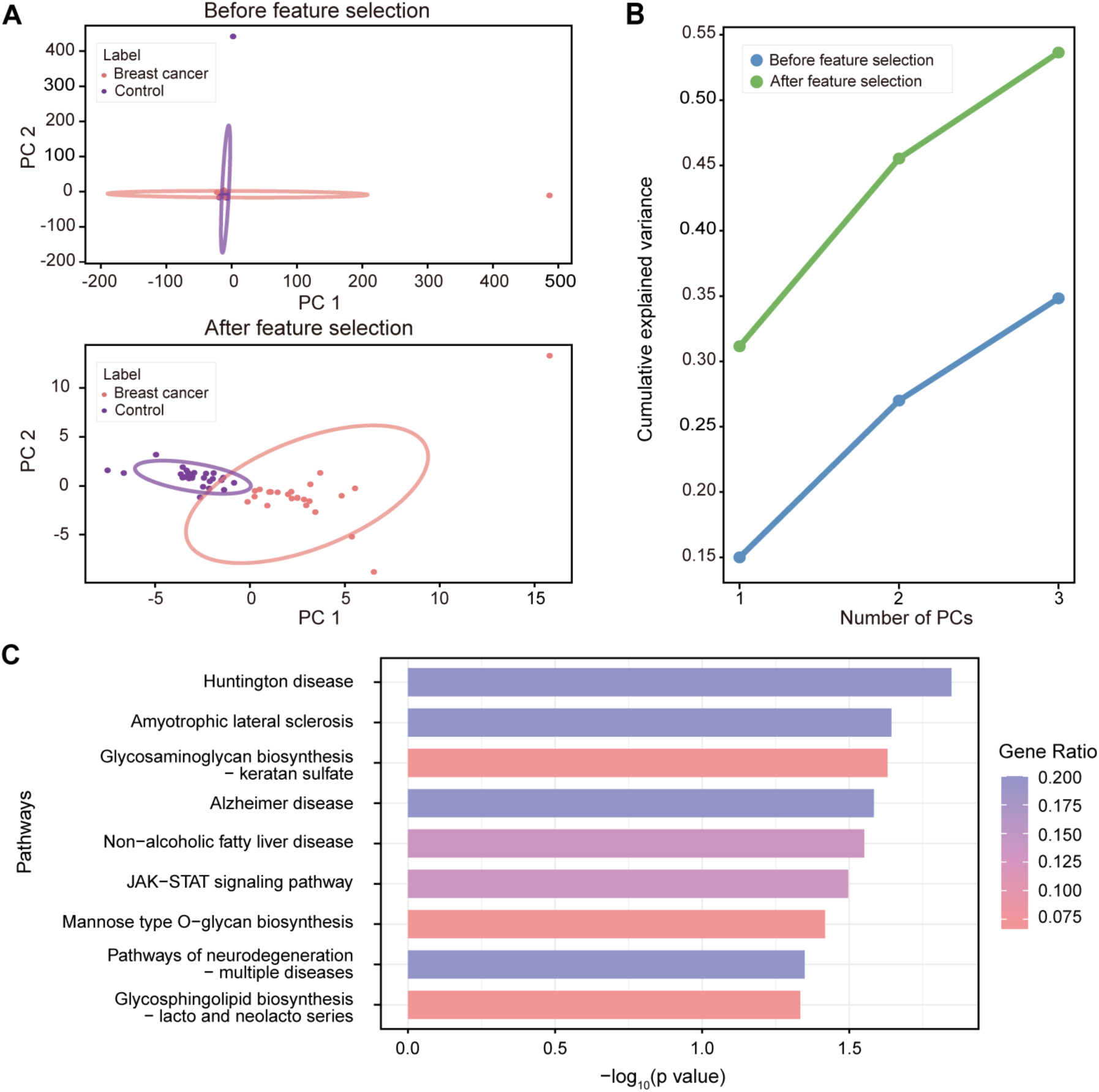
Impact of feature selection and functional relevance. **(A)** Principal Component Analysis (PCA) plots of the TSSC feature before and after applying Boruta algorithm for feature selection. **(B)** Comparison of cumulative variance explained by the top three principal components (PCs) for TSSC features before and after Boruta-based feature selection. **(C)** KEGG pathway enrichment analysis of genes associated with TSSC regions selected by the Boruta algorithm. Pathways with a P-value < 0.05 are shown.

### Cancer detection with cfDNAanalyzer

The growing availability of diverse cfDNA features^24–26^ has opened new opportunities for improving binary classification models in cancer detection. However, a standardized and unified pipeline for systematically evaluating and integrating these features has been lacking. cfDNAanalyzer addresses this need by enabling users to build single-modality models using widely adopted machine learning algorithms—including Random Forest, Linear Regression, Support Vector Machines, K-Nearest Neighbors, Gaussian Naive Bayes, and eXtreme Gradient Boosting—while supporting flexible feature selection through a range of established methods. Model performance can be comprehensively assessed using metrics such as area under the receiver operating characteristic curve (AUROC), F1 score, accuracy, precision, and recall. To facilitate fair comparison, we developed a composite scoring system that incorporates all five performance metrics along with computational efficiency indicators, including memory usage and runtime (**Methods**).

When applied to the breast cancer dataset, models built on end motif frequency and diversity (EM) and nucleosome profile (NP) features achieved the highest performance (**Fig. 4A; Supplementary Table 5)**, with average AUROC values of 0.94 and 0.93, respectively (**Fig. 4B**), indicating a strong discriminative capability. Among feature selection methods, Random Forest importance consistently ranked among the top performers (**Supplementary** Fig. 6A), aligning with results from earlier PCA-based evaluations (**Supplementary** Fig. 4A). To further guide multi-feature integration, cfDNAanalyzer allows users to compute pairwise correlations between predicted probabilities across different feature types **(Fig. 4C)**, offering insight into potential redundancy or complementarity. For the breast cancer dataset, combining EM and NP features using model-based integration strategies led to improved performance over individual feature types (**Fig. 4D**), illustrating the toolkit’s utility in guiding effective feature fusion.

**Fig. 4.**
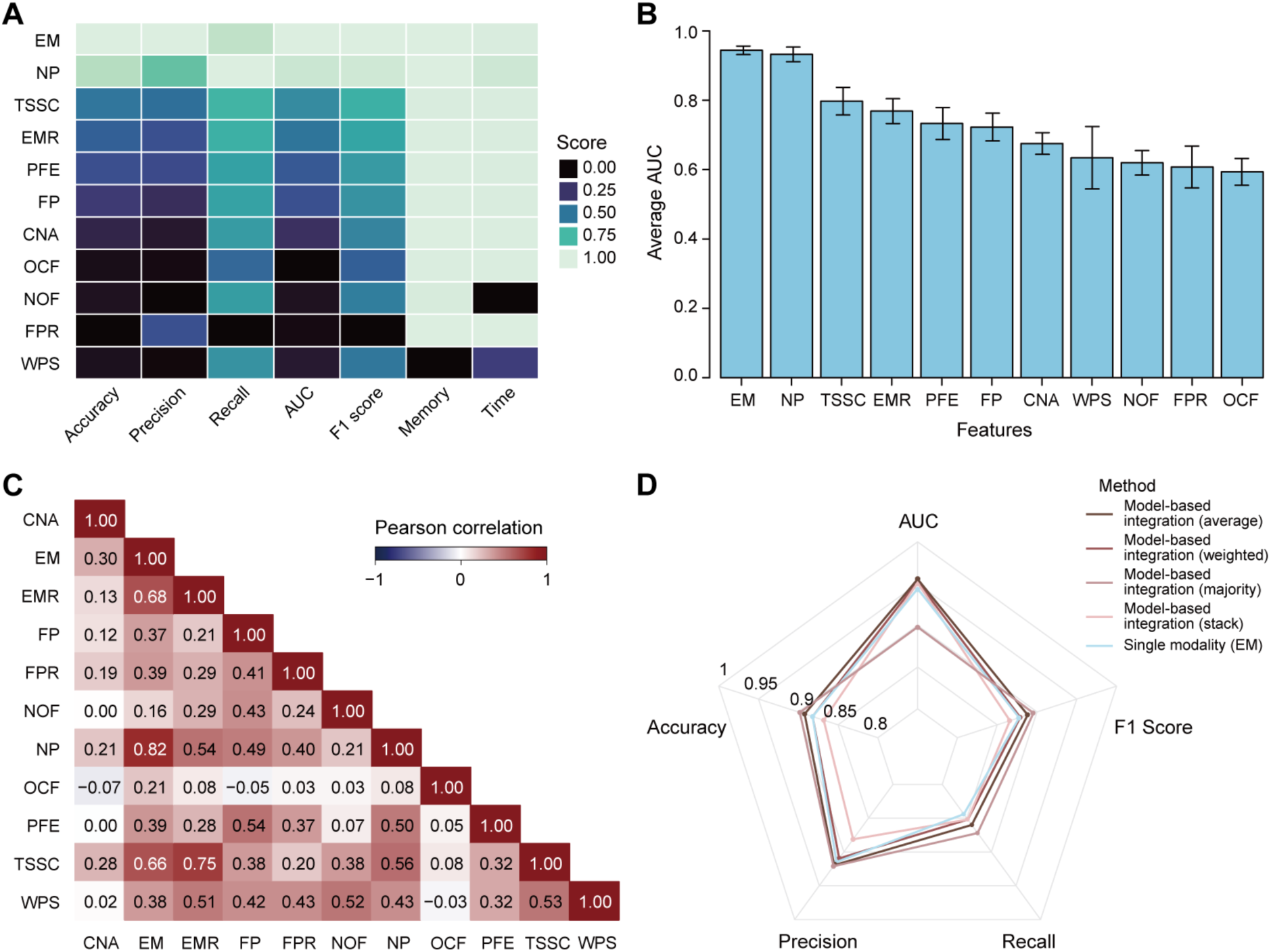
Two-class disease detection performance using cfDNAanalyzer. **(A)** Composite scores for single-modality features, integrating five performance metrics (Accuracy, Precision, Recall, F1-score, AUROC) and two usability metrics (Memory usage, Runtime). **(B)** Bar plots showing AUROCs of 11 single-modality features. Error bars represent standard deviations from 10 repetitions of 5-fold cross-validation. For each feature, only the combination of the feature selection method and classifier with the highest AUROC is shown. **(C)** Pearson correlation matrix of predicted cancer probabilities across all single-modality features, averaged over 10×5-fold CV, highlighting the similarity of prediction outputs. **(D)** Performance comparison of four multi-modality integration strategies against the best-performing single-modality model. Models were built using the top two features identified in **(A)**.

### Cancer classification with cfDNAanalyzer

Beyond binary disease detection, cfDNA analysis plays a crucial role in cancer type classification—an essential step for identifying the tissue of origin and guiding treatment decisions. To demonstrate this functionality, we applied it to a cfDNA WGS dataset comprising samples from lung, breast, and pancreatic cancer patients. The toolkit efficiently handled the entire workflow, from feature extraction to model construction, enabling the development of both single-modality and multi-modality classification models.

Among the single-modality models, the orientation-aware cfDNA fragmentation (OCF) feature achieved the highest overall performance based on our composite scoring system (**Fig. 5A; Supplementary Table 6**), with an average classification accuracy of 0.65 (**Fig. 5B**). In terms of feature selection methods, Fisher’s Score again emerged as a top-performing approach (**Supplementary** Fig. 6B), in line with results from PCA-based evaluations (**Supplementary** Fig. 5A). To assist users in selecting informative features for integration, cfDNAanalyzer also includes tutorials to display the predicted probabilities for each cancer type, helping identify those features that best distinguish among the classes (**Fig. 5C; Supplementary. Fig. 7**). Given the inherent complexity of multi-class cancer classification, we selected five features based on their individual classification accuracy for integration. The resulting integrated model demonstrated improved performance over the best single-modality model in at least one evaluation metric (**Fig. 5D**), highlighting the value of feature integration for enhancing multi-class classification accuracy.

**Fig. 5.**
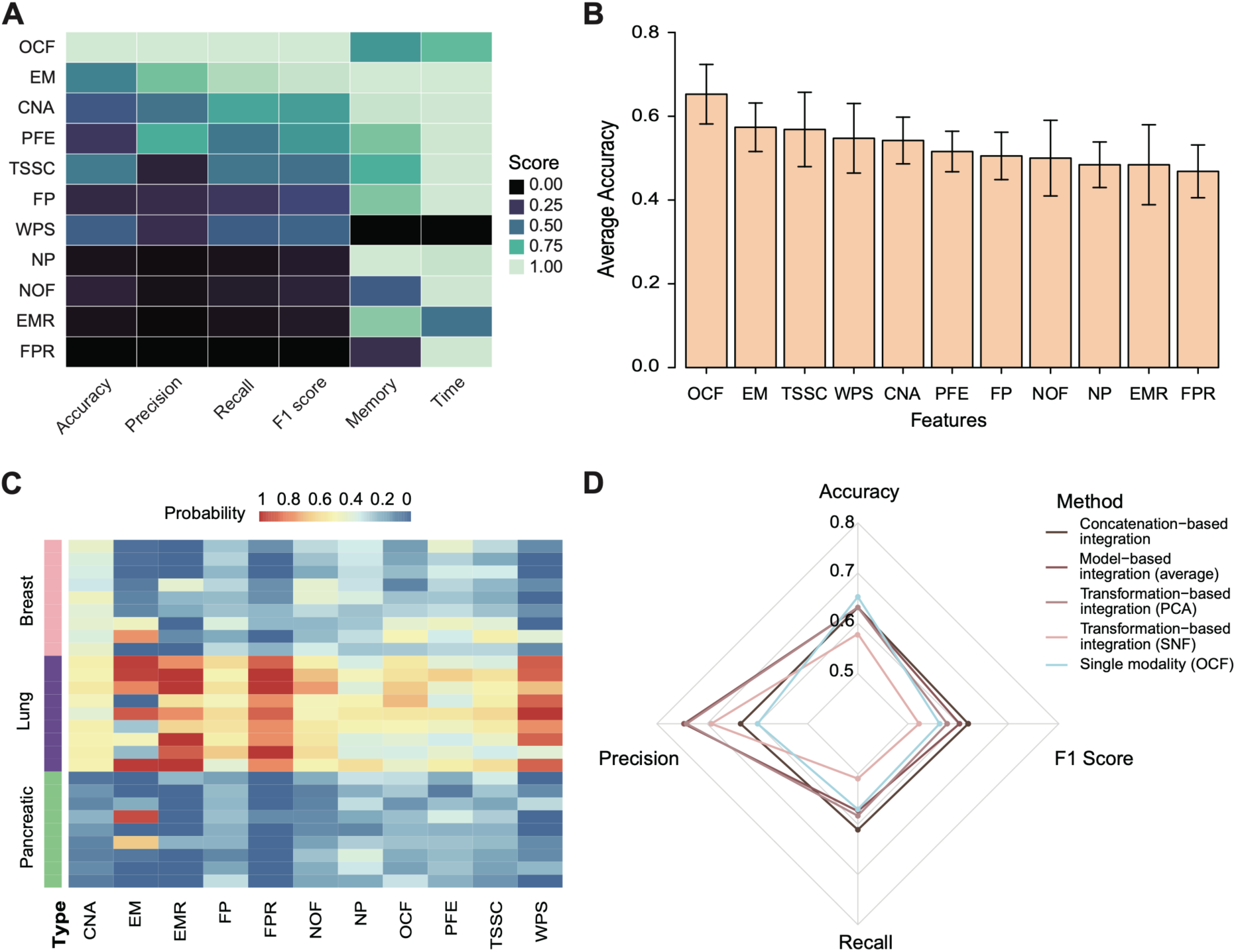
Multi-class disease classification performance using cfDNAanalyzer. **(A)** Composite scores for single-modality features, integrating four performance metrics (Accuracy, Precision, Recall, F1-score) and two usability metrics (Memory usage, Runtime). **(B)** Accuracy comparison of single-modality features for multi-class classification (breast, lung, and pancreatic cancers). Error bars represent standard deviations from 10 repetitions of 5-fold cross-validation. For each feature, only the combination of the feature selection method and classifier with the highest accuracy is shown. **(C)** Heatmap of predicted class probabilities for lung cancer samples across all 11 single-modality features, averaged over 10×5-fold CV. Each row represents a sample; columns represent predicted probabilities for breast, lung, and pancreatic cancers, respectively. **(D)** Performance comparison of four multi-modality integration strategies versus the top single-modality model. Models were built using the five top-ranked features from (**A**).

## Discussion

We present cfDNAanalyzer, a comprehensive and modular toolkit designed to streamline the analysis of cell-free DNA (cfDNA) genomic sequencing data. This versatile platform supports the extraction and interpretation of both genomic and fragmentomic features from single-end and paired-end sequencing data, offering researchers a robust framework for exploring the biological complexity and diagnostic potential of cfDNA. More than just a feature extraction tool, cfDNAanalyzer integrates advanced machine learning functionalities, enabling disease detection and classification through both single-modality and multi-modality models. A key strength of cfDNAanalyzer lies in its wide array of analytical capabilities. The toolkit implements 29 feature processing and selection methods, supporting six widely used machine learning algorithms. This facilitates flexible model construction and comparative evaluation across multiple analysis strategies. Its demonstrated high accuracy in cancer detection and tissue-of-origin classification highlights its potential clinical utility, particularly in the context of non-invasive diagnostics.

What sets cfDNAanalyzer apart is its all-in-one command-line interface, which unifies the entire analysis pipeline—from raw data input to predictive model output—under a single command with customizable parameters. This seamless design allows researchers to efficiently adapt to ongoing innovations in cfDNA feature extraction, as new modules can be incorporated without disrupting the core workflow. This modular and extensible architecture ensures that the toolkit remains future-proof and responsive to the rapidly evolving landscape of cfDNA research and liquid biopsy technologies^27,28^.

Through a range of illustrative case studies, we demonstrate the utility of cfDNAanalyzer in multiple key scenarios: feature extraction and selection, comparative feature exploration, and cancer classification using real-world cfDNA datasets. While currently tailored for cfDNA whole-genome sequencing (WGS) data, cfDNAanalyzer is also compatible with emerging sequencing platforms optimized for fragmentomics analysis, including cfTAPS and cfDNA EM-seq^29,30^, underscoring its adaptability across evolving technological applications.

Importantly, cfDNAanalyzer is designed with broad accessibility in mind. It is freely available and accompanied by extensive documentation and tutorials, guiding users through the whole analytical process—from raw data preprocessing to model training and evaluation. Its intuitive interface lowers the barrier for users with limited computational backgrounds, such as experimental biologists and clinicians, while still offering the flexibility and scalability required by experienced bioinformaticians and data scientists. This inclusive design makes cfDNAanalyzer suitable not only for academic research labs but also for clinical research units and core facilities in biotech and biomedical companies seeking rapid and reproducible analysis of cfDNA data.

In summary, cfDNAanalyzer addresses a critical gap in cfDNA data analysis by offering an integrated, modular, and user-friendly solution. Its powerful combination of feature engineering, machine learning, and flexible design positions it as a valuable tool for advancing cfDNA-based diagnostics, with broad applicability across research, translational, and clinical domains. Future updates will focus on expanding compatibility with additional cfDNA-derived data types and incorporating deep learning modules to further enhance predictive performance^31,32^.

## Methods

### Implementation details

cfDNAanalyzer is a command-line toolkit designed for the analysis of cell-free DNA. At its core is the shell script, *cfDNAanalyzer.sh*, which accepts input files and various parameters. This script orchestrates the execution of integrated scripts to extract features from input Binary Sequence Alignment Map (BAM) files and process these features for downstream analysis. The toolkit offers general parameters as well as many feature-specific options, allowing users to customize the feature extraction and analysis processes according to their needs.

#### Feature extraction

cfDNAanalyzer is designed to extract a comprehensive set of cfDNA-derived features, which can be broadly categorized into three groups:

1. **Genome-wide features**: These features are extracted either across the entire genome at fixed intervals or in aggregate form. Examples include Copy Number Alterations (CNA) and Fragmentation Profiles (FP) computed over tiled genomic bins, as well as End Motif frequency and diversity (EM), which are assessed globally without spatial constraints.
2. **Region-specific features**: These features are calculated within user-defined genomic regions provided in BED format. This category includes Nucleosome Occupancy and Fuzziness (NOF), Windowed Protection Score (WPS), Orientation-aware cfDNA Fragmentation (OCF), End Motif frequency and diversity for Regions (EMR), Fragmentation Profiles for Regions (FPR), and Nucleosome Profiles (NP), enabling targeted analysis in biologically or clinically relevant loci.
3. **TSS-based features**: These are computed in the vicinity of TSSs to capture promoter-level fragmentomic signals. They include Promoter Fragmentation Entropy (PFE), which quantifies the diversity of fragment lengths within ±1 kb of TSSs, and TSS Coverage (TSSC), representing sequencing read coverage in promoter-flanking regions.

#### Feature processing and selection

cfDNAanalyzer first formatted the feature matrix and removed missing values to ensure data integrity. Then, each matrix can be standardized using one of the four methods: Z-score normalization (Zscore), Min-Max Scaling (MMS), Robust Scaling (RS), or Quantile Transformation (QT). The choice of method depended on the data’s distribution: Z-Score normalization centered features to a mean of zero and unit variance; Min-Max Scaling rescaled values to a range between 0 and 1; Robust Scaling used medians and interquartile ranges to reduce the impact of outliers; Quantile Transformation mapped feature values to a uniform distribution.

Feature selection could be conducted using a comprehensive suite of methodologies, including filter-based, wrapper-based, embedded-based, and hybrid approaches:

- Filter-based methods assessed features using statistical and correlation-based measures, including Information Gain (IG), Chi-square Test (CHI), Fisher’s Score (FS), Fast Correlation-Based Filter (FCBF), Permutation Importance (PI), Correlation Coefficient (CC), Low Variance Filter (LVF), Mean Absolute Difference (MAD), Dispersion Ratio (DR), Mutual Information (MI), ReliefF (RLF), SURF, MultiSURF (MSURF), and TuRF (TRF). These techniques systematically ranked features based on intrinsic data properties without relying on a predictive model.
- Wrapper-based methods iteratively refined feature subsets by assessing model performance, using techniques such as 1) Boruta (BOR), a random forest-based all-relevant feature selection approach; 2) Stepwise selection, including Forward Feature Selection (FFS) and Backward Feature Selection (BFS); and 3) Exhaustive search, including Exhaustive Feature Selection (EFS), and Recursive Feature Selection (RFS).
- Embedded methods integrated feature selection within the model training process, including LASSO, Ridge Regression (RIDGE), ElasticNet (ELASTICNET), and Random Forest Importance (RF).
- Hybrid approaches combine filter, wrapper, and embedded techniques to leverage their complementary strengths. For example, in the Filter–Wrapper (FW) strategy, a filter method first ranks and reduces the feature set, followed by wrapper-based selection. In the Filter– Embedded (FE) approach, filtering narrows the feature space before applying an embedded model. These hybrid strategies often yield more robust and reliable feature subsets.

#### Construction of machine learning models

For model construction, cfDNAanalyzer implemented both single-modality and multi-modality integration approaches to maximize the predictive power of cfDNA-derived features. For single-modality models, cfDNAanalyzer includes commonly used algorithms, such as Support Vector Machine (SVM), Random Forest (RF), eXtreme Gradient Boosting (XGBoost), K-Nearest Neighbors (KNN), Gaussian Naive Bayes (GaussianNB), and Logistic Regression. These algorithms were selected for their complementary strengths in handling high-dimensional and potentially correlated biological data.

To leverage complementary signals across different feature modalities, cfDNAanalyzer implemented three multi-modality integration strategies^33^:

- Concatenation-based integration: Selected features from each modality were merged into a single composite feature vector per sample, enabling classifiers to learn from the unified representation.
- Model-based integration: Separate classifiers were trained on the features of each modality. At the prediction stage, their outputs were combined using ensemble techniques, including average voting, weighted voting, majority voting, and a stacking ensemble strategy, to generate a final consensus prediction.
- Transformation-based integration: Each modality’s feature set was first projected into a lower-dimensional embedding space using techniques such as Principal Component Analysis (PCA) and kernel-based dimensionality reduction (e.g., linear, polynomial, radial basis function (RBF), and sigmoid kernels), as well as Similarity Network Fusion (SNF). The resulting embeddings were then concatenated for joint classification, capturing latent relationships across modalities.

To ensure robust evaluation of model generalizability, cfDNAanalyzer applied cross-validation strategies, including K-fold and leave-one-out (LOO) validation, and provided five performance metrics—AUROC, accuracy, precision, recall, and F1 score—to evaluate the model performance:

- AUROC: Measures the model’s ability to discriminate between classes across all classification thresholds, with higher values indicating better performance.
- Accuracy: The proportion of correct predictions (both true positives and true negatives) out of all predictions.
- Precision: The fraction of true positive predictions among all instances predicted as positive.
- Recall: The proportion of actual positives (e.g., diseased samples) correctly identified by the model.
- F1-score: The harmonic mean of precision and recall, balancing both metrics into a single value.

### Processing of data for the figures

#### Datasets

The datasets used in this study are publicly available on Gene Expression Omnibus (GEO) under the accession number GSE71378 (multi-cancer dataset)^2^ and Zenodo with the URL https://doi.org/10.5281/zenodo.6914806 (breast cancer dataset)^34^. Detailed sample information is provided in **Supplementary Table 4**. For the multi-cancer dataset, raw sequencing reads were filtered to retain only those longer than 35 bp and aligned to the human reference genome (GRCh37) using Bowtie2 (v2.2.7)^35^ with the parameters *--mm --no-mixed -X 1000*. The resulting BAM files were further filtered using SAMtools (v1.6)^36^ with the options *-q 30 -F 1804 -f 3* to retain only high-quality, properly mapped reads. For the breast cancer dataset, downloaded fragment files were converted into BAM files using the Fragmentstein tool (v2023.4a)^37^ and subsequently filtered using SAMtools (v1.6) with the parameters -*q 10 -F 1796* to ensure read quality.

#### Feature processing, selection, and evaluation

Leveraging the all-in-one functionality of cfDNAanalyzer, we conducted feature extraction, filtering, and selection from the processed BAM files using a single command: *sh cfDNAanalyzer.sh -s pair -g hg19 --standardM Zscore --noML -F CNA,EM,FP,NOF,NP,WPS,OCF,EMR,FPR,PFE,TSSC --filterMethod ’IG CHI FS FCBF LVF MAD DR MI RLF SURF MSURF ’ --filterNum 50 --wrapperMethod ’BOR’ --wrapperNum 50 -- embeddedMethod ’LASSO RIDGE ELASTICNET RF’ --embeddedNum 50*. Input files and output directories were set accordingly. For region-specific features, we used ±1000 bp surrounding all refSeq genes’ TSS. This command extracted all supported cfDNA features, removed missing values, applied Z-score standardization, and performed feature selection using available filter, wrapper, and embedded methods implemented in cfDNAanalyzer.

For the comparison of PFE between different gene sets, we downloaded expression data of Lung Squamous Cell Carcinoma (LUSC, n=552) and Breast Invasive Carcinoma (BRCA, n=1,226) from the UCSC Xena database (URLs: https://gdc-hub.s3.us-east-1.amazonaws.com/download/TCGA-LUSC.star_tpm.tsv.gz and https://gdc-hub.s3.us-east-1.amazonaws.com/download/TCGA-BRCA.star_tpm.tsv.gz). Gene expression values were originally provided in log2(TPM + 1) format, where TPM refers to transcripts per million. These values were reverse-transformed to raw TPM, and the average TPM across all tumor samples was then computed to represent the tumor expression profile. RNA-seq data for peripheral blood mononuclear cells (PBMCs; n=13) were downloaded from Gene Expression Omnibus (GEO) under the accession number GSE107011^38^. The raw sequencing reads were quality-filtered using TrimGalore (v0.6.10, *-q 25 --illumina -- phred33 --length 35 --stringency 3 –paired*). The remaining reads were aligned to the GRCh38 reference genome using STAR^39^ (v2.7.10b, *--runMode alignReads --readFilesCommand zcat -- quantMode GeneCounts*), employing a gene annotation file consistent with the tumor data. We then normalized the resulting gene counts to TPM and computed the average TPM across all PBMC samples. We limited our analyses to protein-coding genes and removed genes with TPM > 100,000 in LUSC tumors and PBMCs (n=19,962 genes). To identify genes highly expressed in LUSC tumors but lowly expressed in PBMCs, we selected genes with TPM > 40 in LUSC tumors and TPM < 2 in PBMCs (n=459 genes). Conversely, we selected genes lowly expressed in LUSC tumors with TPM < 2 in LUSC tumors and TPM > 40 in PBMCs (n=38 genes). Following the same strategy, we selected genes highly (n=481) and lowly (n=62) expressed in BRCA tumors.

Finally, we performed PCA on each feature type before and after feature selection and calculated the change in cumulative variance explained by the top three principal components to assess dimensionality reduction effectiveness. Functional enrichment analysis of selected features was carried out using ChIPseeker (v1.38.0)^40^, with the KEGG database as the reference^41^.

#### Single-modality model construction and evaluation

For the breast cancer dataset, we constructed single-modality binary classification models to distinguish between breast cancer patients and healthy individuals using the following command: *sh cfDNAanalyzer.sh -F CNA,EM,FP,NOF,NP,WPS,OCF,EMR,FPR,PFE,TSSC -g hg19 -f hg19.fa --labelFile label.txt --filterMethod ’IG CHI FS FCBF LVF MAD DR MI RLF SURF MSURF ’ --filterNum 50 --wrapperMethod ’BOR’ --wrapperNum 50 --embeddedMethod ’LASSO RIDGE ELASTICNET RF’ --embeddedNum 50 --classNum 2 --cvSingle KFold*. For the multi-cancer dataset, which includes cfDNA WGS data from breast, lung and pancreatic cancer patients, we constructed single-modality three-class models using: *sh cfDNAanalyzer.sh -F CNA,EM,FP,NOF,NP,WPS,OCF,EMR,FPR,PFE,TSSC -g hg19 -f hg19.fa --labelFile label.txt -- filterMethod ’IG CHI FS FCBF LVF MAD DR MI RLF SURF MSURF ’ --filterNum 50 -- wrapperMethod ’BOR’ --wrapperNum 50 --embeddedMethod ’LASSO RIDGE ELASTICNET RF’ --embeddedNum 50 --classNum 3 --cvSingle KFold*. Input files and output directories were configured appropriately. For region-specific features, we used ±1000 bp surrounding all refSeq genes’ TSS as input regions. These commands invoked cfDNAanalyzer to apply multiple feature selection strategies and machine learning algorithms to each feature type. Model performance was evaluated using 10 repetitions of 5-fold cross-validation. The following five performance metrics were computed on the test folds: AUROC, accuracy, precision, recall, and F1 score.

#### Composite scoring system for single-modality models

To facilitate robust model selection, we developed a composite scoring system that integrates both performance and usability metrics. The five performance metrics (AUROC, accuracy, precision, recall, and F1 score) were combined with two resource-based metrics: runtime and memory usage. Run time was measured by recording the start and end times of each process. Memory usage was monitored using Python’s tracemalloc module, which captures peak memory allocation during execution. All processes were executed using one thread on a computing server equipped with an Intel(R) Xeon(R) Gold 5218R processor with 40 threads and 424 GB of memory, operating on Ubuntu 22.04 LTS with Python version 3.7.16. To compute the composite score, each metric was first normalized using the min-max scaling method. The final score for each model was calculated as: 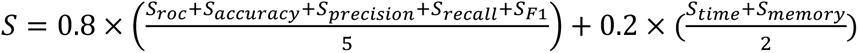, yielding a composite score between 0 and 1, balancing predictive performance and computational efficiency.

#### Multi-modality model construction

Based on the composite scoring results, we constructed multi-modality models using all integration strategies available in cfDNAanalyzer. The top-performing features were selected for integration. For the breast cancer dataset, the top two modalities, EM and NP, were used. For the multi-cancer dataset, the top five modalities—OCF, EM, CNA, PFE, and TSSC—were selected. These selected features were then integrated using cfDNAanalyzer’s concatenation-based, model-based, and transformation-based ensemble strategies to improve classification performance beyond single-modality models.

## Funding

This work was supported by grants from the National Natural Science Foundation of China (Grants No. 82402740), the Natural Science Foundation of Jiangsu Province (Grants No. BK20240783), Soochow University Setup fund, and Jiangsu Distinguished Professor fund to Y.L..

## Conflicts of interest

The authors declare that they have no conflicts of interest.

## Acknowledgments

The authors thank members of Yumei Li’s lab for helpful discussions.

## Supplementary information

Supplementary Figs. 1 to 7, and Tables 1 to 6.

## Notes

### Competing Interest Statement

The authors have declared no competing interest.

